# Phage-resistant bacteria reveal a role for potassium in root colonization

**DOI:** 10.1101/2021.05.12.443821

**Authors:** Elhanan Tzipilevich, Philip N. Benfey

## Abstract

Bacteriophage predation is an important factor in bacterial community dynamics and evolution. Phage-bacteria interaction has mainly been studied in lab cultures, while dynamics in natural habitats, and especially in the plant root niche are underexplored. To better understand this process, we characterized infection of the soil bacterium *B. subtilis* NCBI 3610 by the lytic phage SPO1 during growth in LB medium and compared it to root colonization. Resistance *in vitro* was primarily through modification of the phage receptor. However, this type of resistance reduced the ability to colonize the root. From a line that survived phage infection while retaining the ability to colonize the root, we identified a plant-specific phage resistance mechanism involving potassium (K+) ion influx modulation and enhanced biofilm formation.

## Introduction

Plant roots are associated with diverse bacteria in the soil (Lundberg et al., 2012), which can affect many aspects of plant life including root architecture (Verbon and Liberman, 2016), nutrient acquisition (Hacquard et al., 2015) and disease state (Lugtenberg and Kamilova, 2009; Verbon and Liberman, 2016). The root rhizosphere, i.e. the area close to the root surface, is enriched with specific bacterial taxa in comparison to bulk soil. This unique microbial composition is determined by the soil surrounding the plant root and its pre-existing bacterial diversity (Finkel et al., 2019; Peiffer et al., 2013), plant genotype (Wagner et al., 2016), as well as the interaction with other bacteria (Finkel et al., 2020), and with phage (Koskella and Taylor, 2018). Although a large body of research has been conducted to characterize each of these factors, the whole picture, and especially the role of phage in bacterial root colonization, is far from complete.

Phage are viruses that infect and kill bacteria (Koskella and Brockhurst, 2014; Salmond and Fineran, 2015). As bacterial predators, they influence bacterial community dynamics through elimination of their sensitive hosts. Bacteria, in turn, respond by rapid evolution of phage resistance (Labrie et al., 2010). In addition to influencing bacterial interaction with phage, newly acquired resistance (Koskella and Brockhurst, 2014; Rodriguez-Valera et al., 2009) can also cause changes in colony morphology (Buckling and Rainey, 2002), genome wide mutation rate (Pal et al., 2007) and lateral spread of genetic material (Haaber et al., 2016). In recent years, insights have been gained into how phage-bacteria interactions affect bacterial growth in ocean (Paul and Sullivan, 2005), and mammalian gut ecosystems (Shkoporov and Hill, 2019). However, little is known about how the effects of phage-bacteria interactions in other ecosystems, especially in soil and the root rhizosphere (Pratama and van Elsas, 2018).

To address this question, we utilized *Bacillus subtilis* strain NCIB 3610 (henceforth *B*.*subtilis*) (Branda et al., 2001) and its cognate lytic phage SPO1 (Stewart et al., 2009) to explore their interaction during root colonization of the plant model system *Arabidopsis thaliana. B. subtilis* is a Gram-positive, spore forming bacteria isolated from soil, and is able to colonize plant roots (Chen et al., 2012). Root colonization by *B. subtilis* is mediated by formation of a biofilm on the root (Beauregard et al., 2013). Biofilms are bacterial communities encased in an extracellular matrix. The *B. subtilis* matrix is mainly composed of sugar polymers, encoded by the *eps* operon, and protein fibers encoded by *tapA-sipW-tasA* operon (Vlamakis et al., 2013). *B. subtilis* defective in biofilm formation exhibit severe defects in root colonization (Beauregard et al., 2013). The phage SPO1 is a lytic phage, representing a large and diverse group of Myoviridae bacteriophages, harboring a long contractile tail. SPO1 phage exhibit a complex infection cycle, involving the subversion of the host transcription machinery for its own use (Hoet et al., 1992). SPO1 utilize the wall teichoic acid polymers (WTA) as a receptor to invade *B. subtillis* cells (Habusha et al., 2019). WTA are long sugar polymers that are incorporated into the cell wall and membrane of *B. subtillis* and other Gram-positive bacteria (Brown et al., 2013). SPO1 bind these polymers when they are decorated by glucose moieties (gWTA). Comparison of phage-bacteria evolution upon infection in LB medium and during root colonization revealed that phage infection *in vitro* and i*n planta* exhibits different evolutionary trajectories. Characterization of bacteria resistant to phage infection during root colonization led to the identification of a plant-specific phage resistance mechanism through modulation of potassium (K^+^) ion influx, and enhanced biofilm formation. Furthermore, we show that potassium serves as a stimulator of root colonization among diverse bacilli species.

## Results

### Loss of phage receptor results in a fitness cost for root colonization

To explore phage-bacteria interactions during root colonization we inoculated the roots of *A. thaliana* with *B. subtillis* 3610 bacteria together with SPO1 phage at either high (phage:bacteria 1:1), or low (phage:bacteria 1:10) multiplicity of infection (MOI). Measuring bacterial colonization (CFU colony forming units) after 48 hours, revealed a reduction in root colonization of ∼95% at an MOI of 1, and ∼90% reduction at an MOI of 0.1 (Figure 1A) indicating that SPO1 can efficiently infect and kill its host bacteria during root colonization. Because selection pressure by phage drives bacterial counter adaptation (Buckling and Brockhurst, 2012) we isolated 300 bacteria [screen 1] that survived phage infection on the root and re-streaked them on agar plates containing SPO1 phage. Only 20 *in planta* survivors were resistant to phage infection *in vitro*. In contrast, 100% of 70 bacteria isolated after surviving phage infection in LB medium, became immune to further infection. Twenty randomly selected *in vitro* survivors became phage resistant through loss of the phage receptor, as judged by a lack of conA_488_ staining, a lectin that binds specifically glycosylated WTA (gWTA) (Figure 1B) [see also (Habusha et al., 2019)]. Of the 20 SPO1 resistant bacteria isolated from the plant all but one (m28) had lost their ability to re-colonize the root (Figure 1C). Nineteen of these isolates concomitantly lost their conA_488_ staining (Figure S1A). One of the isolates, m28, was completely phage resistant in-vitro (Figure S1B), but still exhibited faint conA_488_ staining (Figure S1A and S1C), and partial sensitivity when infected on roots (Figure 2A). The correlation between SPO1 sensitivity and root colonization ability suggests that gWTA is important for efficient root colonization. To test this hypothesis, we inoculated roots with *ΔtagE* bacteria, which lack gWTA (Allison et al., 2011), and found a significant reduction in root colonization (Figure 1D-1E). Because biofilm formation has been shown to be necessary for root colonization by *B. subtilis* (Banda et al., 2019), we tested the ability of *ΔtagE* bacteria to form a biofilm *in-vitro. ΔtagE* cells exhibit normal biofilm formation both on agar plates and liquid medium (Figure 1F). Of note, gWTA is required for nasal epithelium colonization by *Staphylococcus aureus* (Winstel et al., 2015). Our results suggest that gWTA in *B. subtilis* is similarly important for plant surface adhesion, irrespective of biofilm formation.

**Figure 1.**
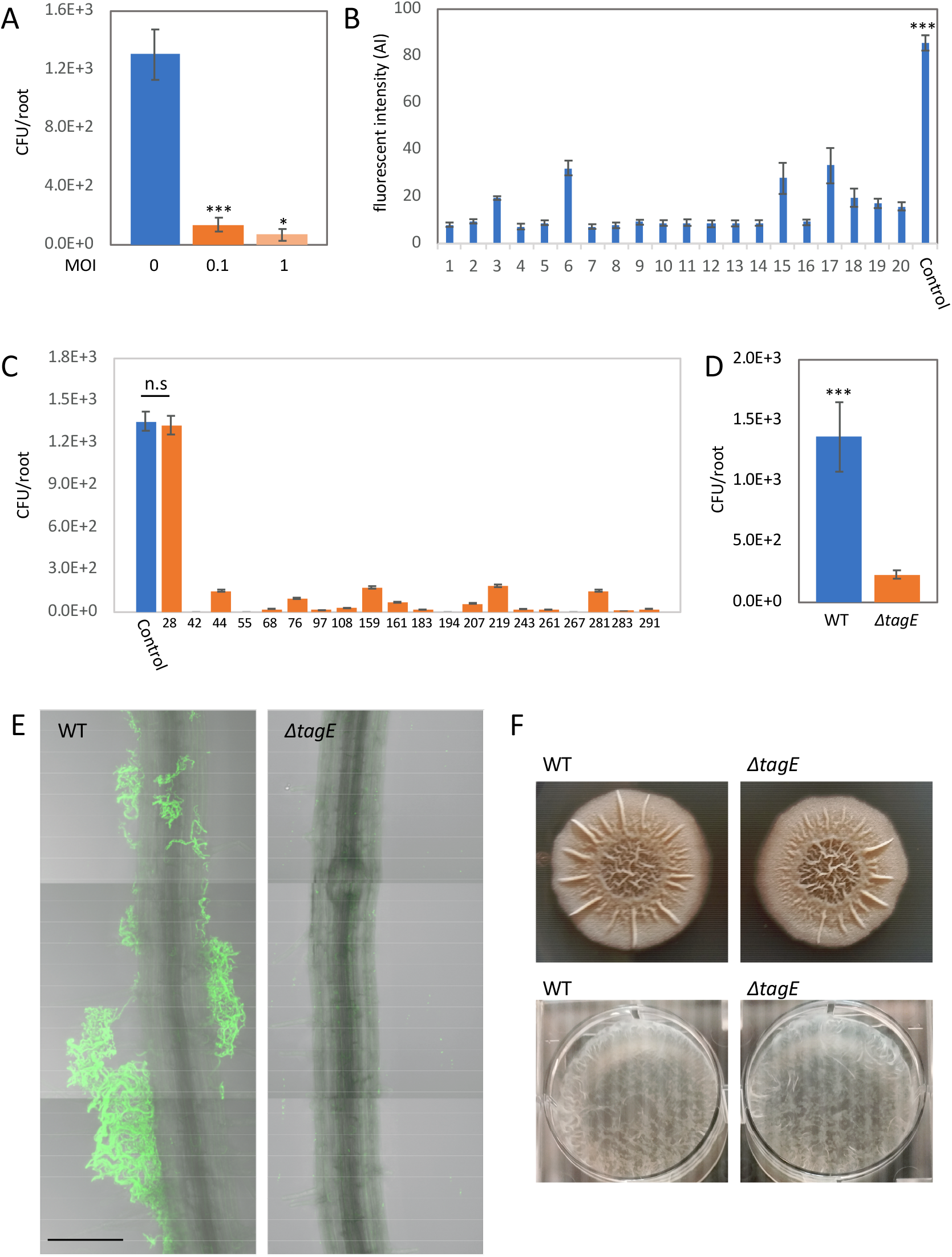
SPO1 phage receptor is necessary for bacterial root adhesion. (**A**) Seedlings were inoculated with *B. subtilis* 3610 together with SPO1 phage, at either high (phage : bacteria 1:1), low (phages : bacteria 1:10) multiplicity of infection (MOI), or no phage (MOI = 0) for 48 hrs, on agar plates, and the number of colonizing bacteria was counted. Shown are averages and SD of 2 independent experiments with n ≥ 3 for each, * = *P <* 0.05, *** = *P <* 0.005. (**B**) 20 randomly selected bacterial colonies that survived SPO1 infection were stained with conA_488_ and observed under a compound microscope. Shown are average and SD fluorescent intensity values of 10 bacteria from each of the colonies. (**C**) 20 isolated SPO1 resistant bacteria were inoculated on seedlings, grown for 48 hrs and the number of colonizing bacteria counted. Shown are averages and SD with n ≥ 3, *** = *P <* 0.005. (**D**) Seedlings were inoculated with either WT or *ΔtagE* bacteria for 48 hrs, and the number of colonizing bacteria was counted. Shown are averages and SD of 2 independent experiments with n ≥ 3 for each, *** = *P <* 0.005. (**E**) Seedlings were inoculated with either WT or *ΔtagE* bacteria expressing GFP (amyE::P_*rrnE*_-gfp) for 48 hrs on agar plates. Shown are representative overlay 200x confocal images of DIC (root) and GFP fluorescence (bacteria). Scale bar, 50μm. (**F**) WT and *ΔtagE* bacteria were inoculated into MSgg media on agar plates (upper) or liquid media (lower). Shown are representative biofilm images.

**Figure 2.**
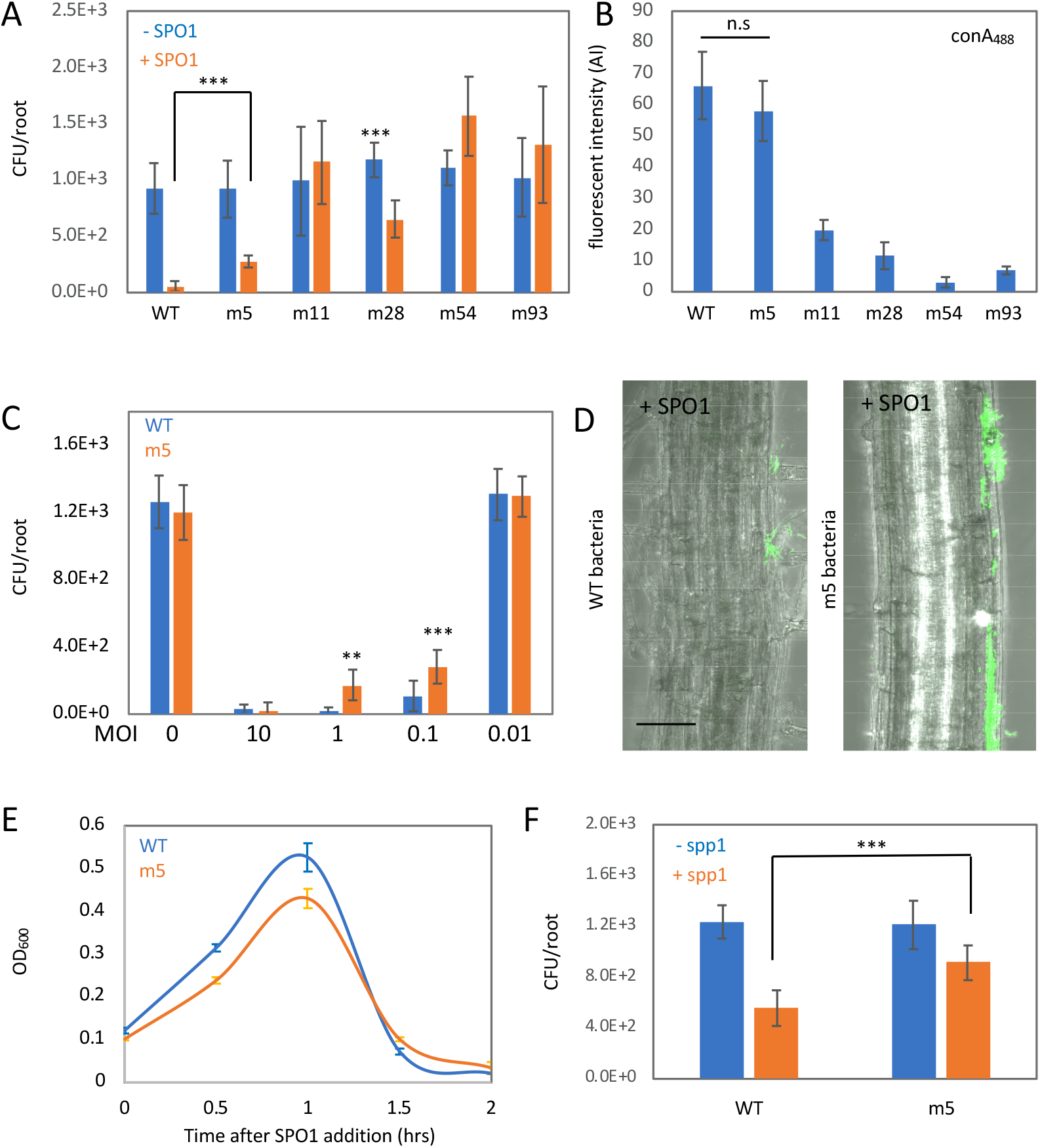
A root-specific SPO1 resistance mechanism. (**A**) Seedlings were inoculated with the indicated bacterial strains, with or without SPO1 addition, for 48 hrs on agar plates, and the number of colonizing bacteria was counted. Shown are averages and SD of 2 independent experiments with n ≥ 3 for each, ** = *P <* 0.01. (**B**) Bacterial strains were stained with conA_488_ and observed under the microscope. Shown are average and SD of fluorescence intensity values of 10 bacteria from each colony. (**C**) Seedlings were inoculated with either WT or m5 bacteria, with the indicated phage MOI, for 48 hrs on agar plates, and the number of colonizing bacteria counted. Shown are averages and SD of 2 independent experiments with n ≥ 3 for each, *** = *P <* 0.005. (**D**) Seedlings were inoculated with either WT or m5 bacteria expressing GFP (amyE::P_*rrnE*_-gfp), in the presence of SPO1, for 48 hrs on agar plates. Shown are representative overlay 200x confocal images of DIC (root) and GFP fluorescence (bacteria). Scale bars 20μm. (**E**) WT and m5 bacteria were infected with SPO1 in LB media, and OD_600_ was followed. Shown are averages and SD, n = 3. (**F**) Seedlings were inoculated with either WT or m5 bacteria, with or without spp1 phages for 48 hrs on agar plates, and the number of colonizing bacteria was counted. Shown are averages and SD of 2 independent experiments with n ≥ 3 for each, *** = *P <* 0.005.

### Bacteria evolve a root-specific phage resistance mechanism

Our results indicate that loss of the phage receptor results in a fitness cost to bacteria during root colonization. To identify alternative pathways utilized by bacteria to resist phage infection during root colonization, we isolated bacteria that survived phage infection on the root (from screen 1) and infected them with SPO1 during root colonization (screen 2). Most of the bacteria exhibited phage sensitivity similar to the parental bacteria. However, of 100 bacterial strains tested, we found 4 that survived SPO1 infection on roots better than WT cells (Figure 2A). To explore the mechanism of phage survival, we sequenced the genomes of m5, m11, m54 and m93, the 4 bacteria isolated during screen 2 along with m28 isolated from screen 1. Table 1 presents the mutations in each of the bacteria.

**Table 1.**

| Bacterial strain | Mutation |
| --- | --- |
| m5 | Missense mutation in KtrC<br>P189T |
| m11 | Mutations in YwbO promoter |
| m28 | Missense mutation GtaB Y170S |
| m54 | Missense mutations in<br>YwrK T334S and CdaA F97L |
| m93 | Frame shift in tagE T1013<br>leading to premature stop<br>codon |

M28 and m93 harbor mutations in gtaB and tagE, genes involved in the WTA glycosylation pathway (figure 2B and figure S1C) (Brown et al., 2013). Interestingly, m11 and m54 don’t have mutations in genes previously implicated in WTA glycosylation, but nonetheless exhibited complete (m54) and partial (m11) loss of conA_488_ staining (figure 2B and figure S1C). m11, m54 and m93 exhibited phage resistance *in vitro* (figure S1B).

One mutant, m5, exhibited enhanced phage resistance when infected in-planta at medium MOI (1-0.1) (Figure 2C-2D), phage sensitivity *in vitro*, and conA_488_ staining similar to WT (Figure 2B,2E and figure S2B) indicating that it harbors a root-specific phage resistance mechanism. The m5 mutant also exhibits enhanced resistance to infection by SPP1, a lytic phage from a different bacteriophage family (siphoviridae), (Figure 2F), indicating that its resistance mechanism is not phage-family specific. M5 harbors a single point mutation in the *ktrC* gene, encoding a low affinity potassium channel. This missense mutation changes Proline 189 into Threonine (KtrC P189T) and resides in a conserved RCK domain, known to bind the regulatory molecule ci-d-AMP (Corrigan et al., 2013; Rocha et al., 2019). ci-d-AMP negatively regulates potassium uptake (Gundlach et al., 2019). We hypothesized that KtrC P189T is a gain of function mutation, affecting the regulation of potassium uptake.

### Potassium enhances root colonization and phage tolerance through modulation of biofilm formation

Several potassium channels are encoded in the *B. subtillis* genome: *ktrAB* and *kimA* encode high affinity channels (Gundlach et al., 2017; Holtmann et al., 2003) while the *ktrCD* operon encodes a low affinity channel (Holtmann et al., 2003). To understand the effect of the KtrC P189T mutation and potassium uptake on bacterial root colonization, we examined the colonization efficiency of bacteria lacking these channels. While Δ*ktrC* and Δ*kimA* colonize the root with similar efficiency to WT bacteria (Figure 3A), Δ*ktrA* bacteria exhibited reduced root colonization (Figure 3A). The KtrC P189T mutation is able to restore the colonization of Δ*ktrA* bacteria (Figure 3A), suggesting that this mutation enhances potassium uptake through the KtrC channel. Growing plants on media with different levels of potassium revealed a positive correlation between potassium concentration and root colonization by WT bacteria (Figure 3B), with the effect plateauing at 10mM. Similarly, root colonization on 0.25 MS plates, with addition of 5mM potassium (KCl) (equal to total of ∼10mM) indicated a positive effect of potassium on root colonization (Figure 3C). Stimulation of root colonization was specific to potassium ions, as neither sodium nor nitrogen (Fig S2A) had a similar effect. Consistent with the KtrC P189T mutation affecting phage infection through increased potassium uptake, addition of potassium was sufficient to enhance survival of WT cells upon phage infection (Figure 3C-3D).

**Figure 3.**
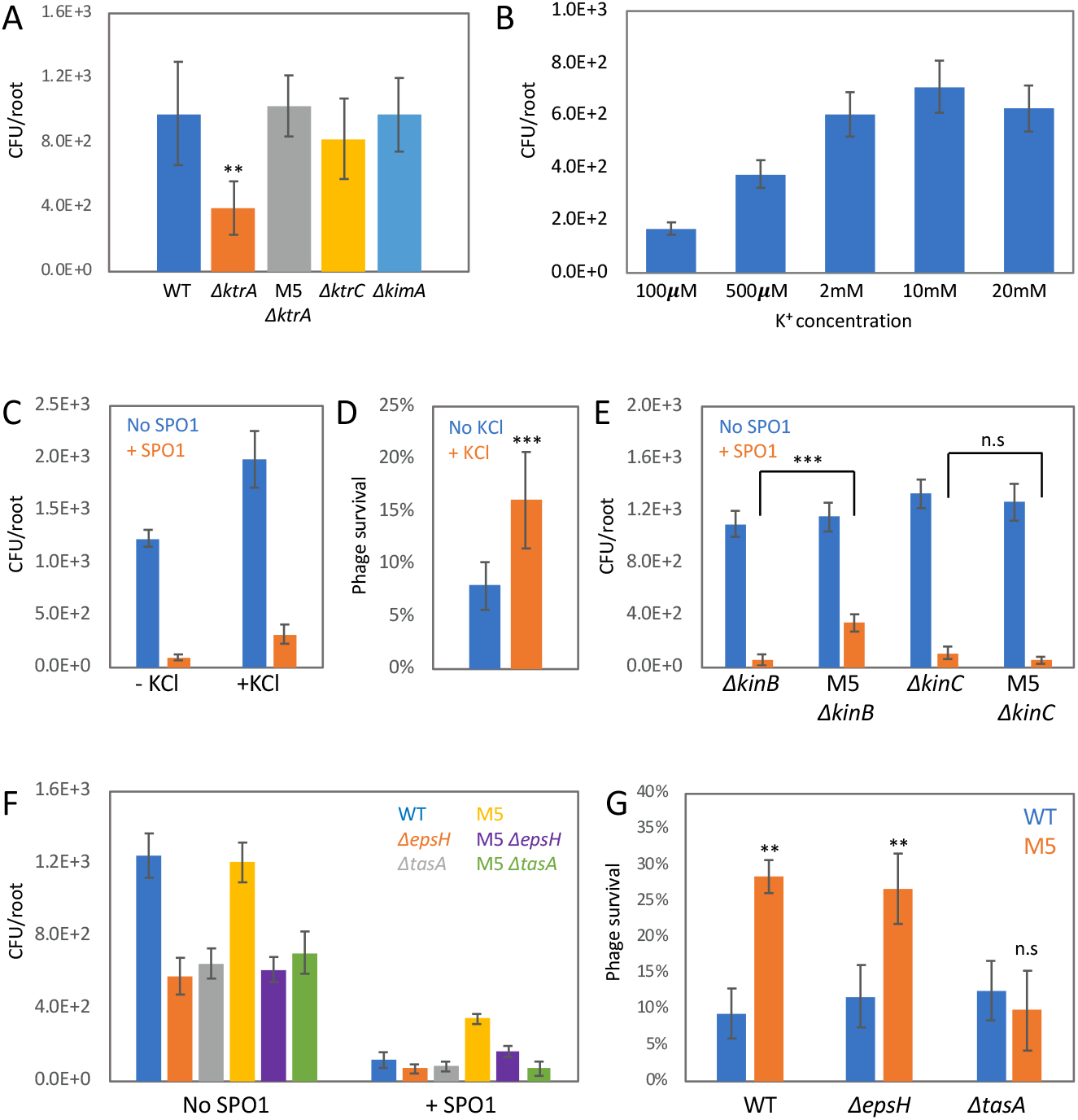
Potassium modulates *B. subtillis* root colonization and phage resistance through biofilm formation. (**A**) Seedlings were inoculated with the indicated bacterial strains for 48 hrs on agar plates, and the number of colonizing bacteria was counted. Shown are averages and SD of 2 independent experiments with n ≥ 3 for each, ** = *P <* 0.01. (**B**) Seedlings were inoculated with WT *B. subtillis* on MS agar plates supplemented with the indicated potassium concentration, for 48 hrs, and the number of colonizing bacteria was counted. Shown are averages and SD of 2 independent experiments with n ≥ 3 for each. (**C-D**) Seedlings were inoculated with WT *B. subtillis*, with or without SPO1 addition, for 48 hrs on 0.25MS agar plates, in the presence or absence of 5mM KCl, and the number of colonizing bacteria was counted. Shown are averages and SD CFU (C), and the percentage of SPO1 survival (D), from 2 independent experiments with n ≥ 3 for each, ** = *P <* 0.01. (**E**) Seedlings were inoculated with the indicated bacterial strains, with or without SPO1 addition, for 48 hrs on agar plates, and the number of colonizing bacteria was counted. Shown are averages and SD of 2 independent experiments with n ≥ 3 for each, ** = *P <* 0.01. (**F-G**) Seedlings were inoculated with the indicated bacterial strains, with or without SPO1 addition, for 48 hrs on agar plates, in the presence or absence of 5mM KCl, and the number of colonizing bacteria was counted. Shown are averages and SD CFU (F), and the percentage of SPO1 survival (G), from 2 independent experiments with n ≥ 3 for each, ** = *P <* 0.01.

It has been previously shown that *B. subtilis* bacteria sense potassium influx through the cell membrane, utilizing KinC and KinB protein kinases to induce biofilm formation genes and sliding motility, respectively, (Grau et al., 2015; Lopez et al., 2009). Both biofilm formation and sliding motility play an important role during root colonization (Bais et al., 2004; Beauregard et al., 2013). We found that m5 Δ*kinC* bacteria, but not m5 Δ*kinB* lost SPO1 resistance (Figure 3E). Addition of potassium to Δ*kinC* bacteria failed to stimulate phage resistance (Figure S2C). KinC activation induces a phosphorylation cascade that culminates in induction of biofilm matrix genes (Lopez et al., 2009). Disruption of the biofilm matrix operons *epsA-O* and *tapA-sipW-tasA* abolished the effect of potassium on root colonization (Figure S2C). Interestingly disruption of the *tapA-sipW-tasA* operon, but not *epsA-O*, reduced the phage resistance phenotype of m5 cells (Figure 3F-3G), suggesting that an increase in protein fiber component is responsible for the phage tolerance effect of increased potassium influx. Thus, our analysis revealed a root-specific phage adaptation mechanism that works through enhanced potassium influx by stimulating biofilm matrix formation.

To further characterize the role of biofilm matrix formation in phage tolerance, we monitored SPO1 infection during biofilm formation *in vitro* (Branda et al., 2001). Measurement of biofilm diameter on MSgg agar plates revealed significant reduction in colony diameter on plates containing SPO1 phage (Figure 4A). Of note, Δ*ktrA* cells, although able to form normal biofilm in the absence of phage, exhibit a significant decrease in biofilm diameter in the presence of phage in comparison to WT infected cells (Figure 4A-4B). Similar to what was observed in plants, the KtrC P189T mutation was able to compensate for the increased sensitivity of Δ*ktrA* cells (Figure 4A-4B). Thus, our *in vitro* analysis, provides further support for the hypothesis that potassium influx modulates biofilm formation to enhance phage resistance. Equivalent results were obtained for pellicle biofilm formation on liquid MSgg medium (Figure S3), where enhanced potassium influx in m5 bacteria, or addition of extra potassium to the medium of WT bacteria, increased the chance of surviving SPO1 infection, while decreased influx, due to the Δ*ktrA* mutation, increased phage sensitivity.

**Figure 4.**
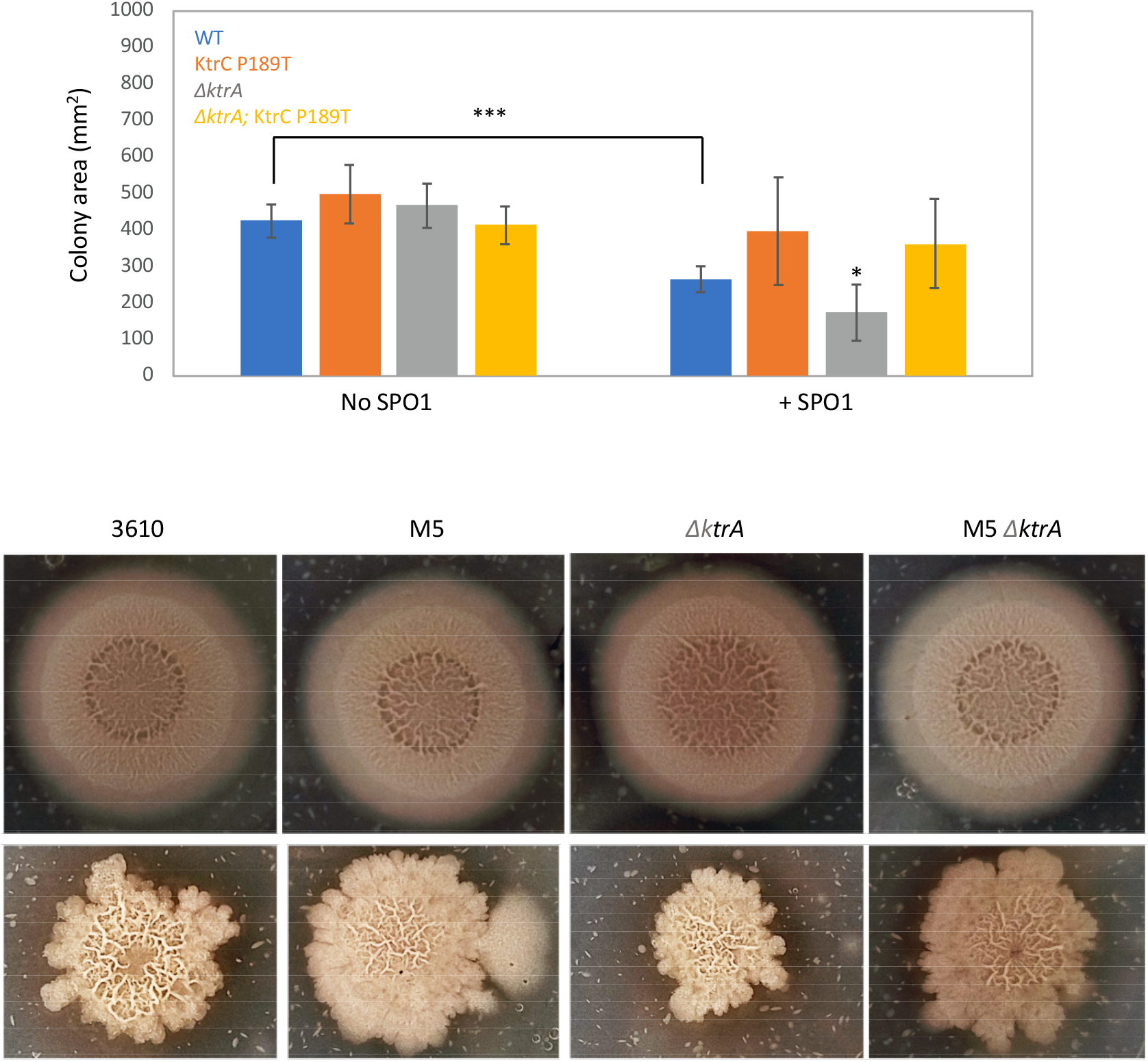
Potassium modulates phage resistance during biofilm formation *in-vitro*. (**A**) The indicated bacterial strain were inoculated onto MSgg agar plates with or without SPO1 addition for 48 hrs, and colony area measured. Shown are averages and SD of n = 6. * = *P <* 0.05. (**B**) Shown are representative images for the experiment described in (A) with (lower panel), or without (upper panel) SPO1 addition.

### Potassium enhances root colonization by diverse bacilli species

To determine if potassium is able to enhance colonization by other bacilli species we analyzed its effect on *B*.*valezensis fzb42* a plant-associated bacterium (Fan et al., 2018), shown to enhance plant health through growth stimulation and pathogen inhibition. We found that potassium enhances root colonization of *B*.*valezensis fzb42* (Figure 5A), and this phenomenon was abolished in *B*.*valezensis fzb42* Δ*kinC* cells (Figure 5B). A similar phenomenon was observed for *B*.*subtilis GB03* and *B*.*pumilus SE34*, two other plant growth promoting bacteria (Figure 5A). Thus, potassium serves as a wide-spread signal, stimulating root colonization by diverse bacilli species. *B*.*valezensis fzb42* inhibits the growth of the plant fungal pathogen *Rhizoctonia solani* (Krober et al., 2014), raising the possibility that simply adding potassium to the growth media could enhance *B*.*valezensis fzb42* colonization and fungal protection. Indeed, potassium enhanced plant survival when inoculated with *Rhizoctonia solani*, in the presence of *B*.*valezensis fzb42* (Figure 5C), and this effect was abolished in Δ*kinC* cells, which are unable to sense potassium influx (Figure 5C).

**Figure 5.**
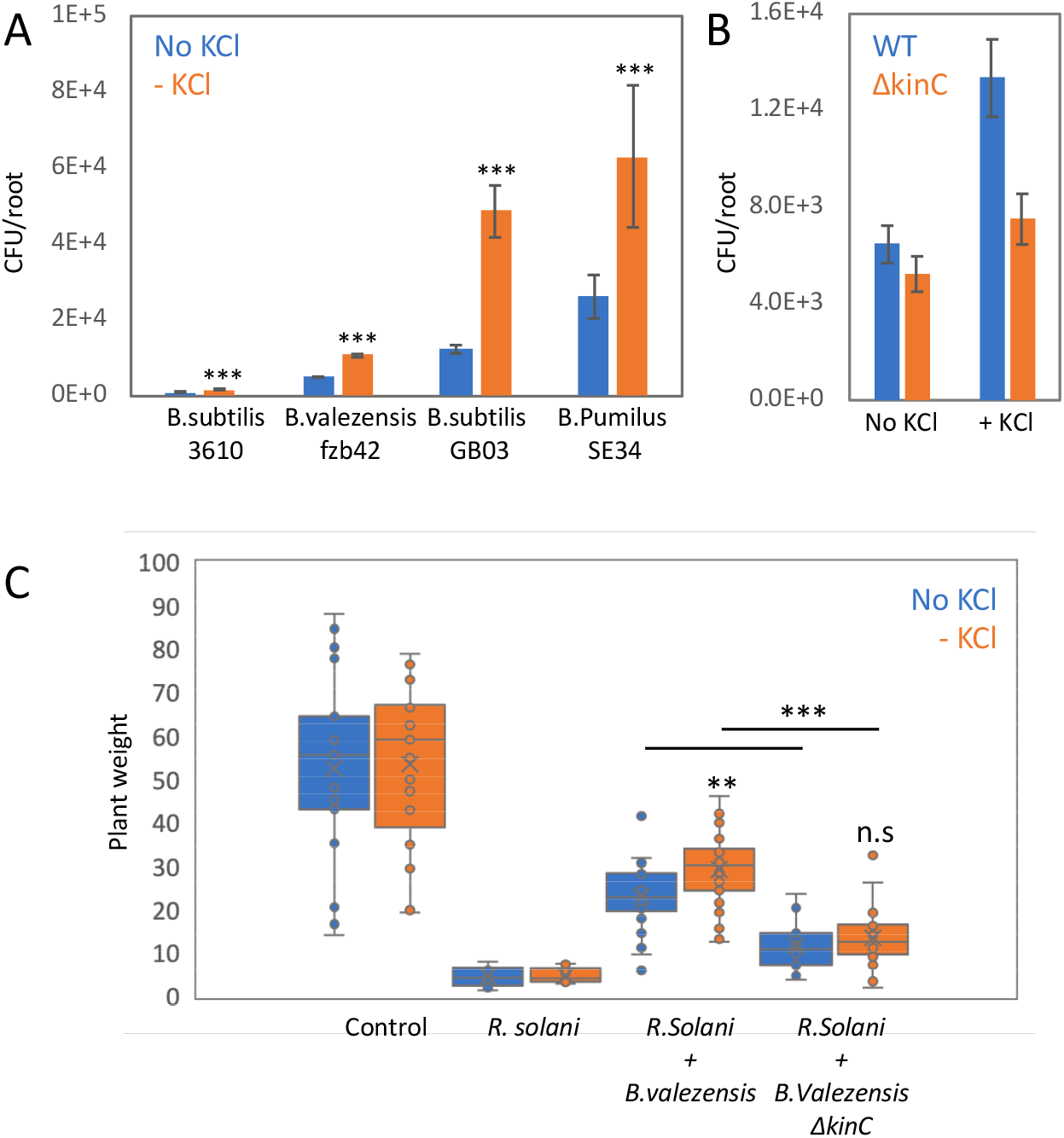
Potassium modulates root colonization by diverse bacilli species. (**A**) Seedlings were inoculated with the indicated bacterial species,for 48 hrs on 0.25MS agar plates, in the presence or absence of 5mM KCl, and the number of colonizing bacteria was counted. Shown are averages and SD of 2 independent experiments with n ≥ 3 for each, ** = *P <* 0.01. (**B**) Seedlings were inoculated with either WT or *ΔkinC B. valezensis fzb42* bacteria, for 48 hrs on 0.25MS agar plates, in the presence or absence of 5mM KCl, and the number of colonizing bacteria was counted. Shown are averages and SD of 2 independent experiments with n ≥ 3 for each, ** = *P <* 0.01. (**C**) Seedlings were inoculated with either WT or *ΔkinC B. valezensis fzb42* bacteria, for 48 hrs on 0.25MS agar plates, in the presence or absence of 5mM KCl. Next, plates were inoculated with *R. solani* and incubated for an additional 7 days and plant weight was measured. Untreated plants (neither bacteria, nor fungi) were used as control. Shown are averages and SD, n ≥ 20. ** = *P <* 0.01, *** = *P <* 0.005.

## Discussion

Phage infection has been extensively characterized in lab cultures. Recent work revealed that phage-bacteria interaction in natural environments can differ significantly from that observed in the lab (Diaz-Munoz and Koskella, 2014) due to the fitness constraints encountered by bacteria in their natural environment. This can impede the evolution of phage receptors, the main resistance pathway observed in the lab (Westra et al., 2015). We found that gWTA, the SPO1 phage receptor is essential for root adhesion, thus receptor modification has a fitness cost in the root environment. Of note, it has been shown that gWTA is essential for adhesion of *Staphylococcus aureus* to the nasal epithelium (Winstel et al., 2015), while gWTA also serves as phage receptor for several *S*.*aureus* infecting phages (Moller et al., 2019). It will be interesting to determine if phage bacteria evolution is constrained in this niche as well.

While evolution of receptor modification is not favored, we found that bacteria in the root adapt to phage infection through evolution of enhanced biofilm formation. Consistent with previous *in-vitro* work (Lopez et al., 2009), we show that during plant colonization, altering potassium efflux through KtrCD channels enhances *B. subtillis* biofilm formation, and thus promotes survival upon phage infection. Our results indicate that protein fiber density, encoded by the *tapA-sipW-tasA* operon, is the main determinant of SPO1 resistance. Interestingly, it has been shown that curli protein fibers are important for T7 resistance in a biofilm of *E. coli* bacteria (Vidakovic et al., 2018). Phage-bacteria interaction in biofilm is an emerging area of research (Hansen et al., 2019), and much remains to be learned about the interaction dynamic in this important bacterial niche. In our screen for *in planta* phage resistant bacteria we found 4 bacteria of 100 with genetically-inherited enhanced phage resistance. This suggests that 96% of the bacteria survived infection in a non-genetic manner. It would be interesting to determine how non-genetic mechanisms enable bacteria to survive phage infection.

Potassium is an essential macronutrient for bacteria and plants. Unlike nitrogen and phosphate, it is not incorporated into cellular macromolecules, but remains a soluble ion (Amtmann and Armengaud, 2009). Potassium is involved in the regulation of many processes, including osmotic adaptation, membrane potential, phloem transport, regulation of metabolic enzymes, and biotic and abiotic stress responses. Potassium, together with nitrogen and phosphate, are the three main ions consumed by plants and are heavily utilized for fertilization in modern agriculture. (Sustr et al., 2019). The effect of nitrogen and phosphorous on root microbial communities has been characterized (Castrillo et al., 2017; Finkel et al., 2019; Leff et al., 2015). However, the role of potassium in bacterial colonization is less well explored. Our analysis suggests that addition of potassium is a simple method to enhance bacilli colonization. Many bacilli species exert positive effects on their plant host by enhancing plant growth, and inhibiting pathogens. An example is our preliminary result that suggests that adding potassium could enhance the ability of *B. valezensis* to protect seedlings from *R. solani* infection.

## Materials and methods

### Bacterial strains and growth conditions

WT *B. subtilis NCBI3610* and AR16 (Rosenberg et al., 2012) (*B. subtilis* PY79) were kindly provided by Prof. Sigal Ben-Yehuda (Hebrew University) *ΔtagE* (BKK35730), *ΔktrA* (BKK31090), *ΔktrC* (BKK14510), *ΔkimA* (BKK04320), (all in *B. subtilis* 168) were purchased from the Bacillus genetic stock center (http://www.bgsc.org/). Genomic DNA was extracted from AR16 and mutant strains using Wizard Genomic DNA Purification Kit (Promega) and transformed into *B. subtilis NCBI3610*. The media and growth conditions used for DNA transformation of *B. subtilis NCBI3610* were described in http://2013.igem.org/Team:Groningen/protocols/Transformation. The bacteria were cultivated routinely on Luria broth (LB) medium. When needed, the medium was solidified with 1.5% agar. For biofilm formation, bacteria grown overnight were inoculated into MSgg medium and incubated without shaking for 2 days at 30° as described in (Branda et al., 2001). This medium was also solidified with 1.5% agar.

### Phage strains and infection conditions

SPO1 (1P4) and SPP1 (1P7) phage were purchased from the Bacillus genetic stock center. Phage lysate was routinely prepared by adding approximately 10^9^ phage to mid-log cells grown in LB medium supplemented with 10mM MgSO_4_, until the culture was completely cleared. The lysate was then filtered through a 0.22µm Millipore filter. For lysis dynamics in LB medium (Figure 2E and Figure S1B), approximately 10^9^ phage were added to mid-log growing cells at multiplicity of infection MOI=1, and OD_600_ was monitored at 30 min intervals. For SPO1 infection in pellicle biofilm (Figure S3), bacteria grown overnight were inoculated into MSgg medium, and phage added to MOI=1. Bacteria were incubated without shaking for 2 days at 30°, and the number of cultures with surviving cells was counted by eye. For SPO1 infection in solid biofilm (Figure 4) bacteria grown overnight were inoculated into MSgg medium supplemented with 10^5^ PFU (plaque forming units), solidified with 1.6% agar. A spot of 5μL from the bacterial culture was inoculated onto the plate and incubated for 2 days at 30°. Plates were scanned and colony diameter determined using ImageJ

### Monitoring bacterial growth and phage infection on plant roots

Plants were grown on 0.5 MS media containing 1.1 gr Murashige and Skoog basal salts (in 500 ml ddH2O), 1% sucrose, 1% agar and 5 ml (in 500 ml ddH_2_O) MES (50 gr/l, titrated to pH=5.8 with NaOH). Plants were stratified for 2 d in a 4°C dark room and grown vertically for 4-10 days under long-day light conditions. Bacteria from fresh colonies were grown in LB medium to an OD_600_ = 1.0 and then diluted 1:100 in PBSx1 for CFU measurements and microscopy, yielding approximately 1×10^6^. Phage were added to the PBSx1 at MOI = 0.1, unless otherwise indicated. Six-day old seedlings were transferred onto square Petri dishes containing 0.5 MS but without sucrose. 2μL of bacterial dilution were deposited immediately above the root tip and left to dry for 2 min. The square plates were kept in a vertical position during the incubation time at 22°C under long-day light conditions (16 h light/8 h darkness) in a plant growth chamber. For bacterial CFU counting and microscopy, plants were incubated with bacteria for 48hrs. Then the inoculated plant roots were cut and washed three times in sterile water. For CFU counting the seedlings were transferred to a tube with 1 ml of PBSx1 and vortexed vigorously for 20 seconds, then a serial dilution was plated on LB plates.

### Microscopy

Roots were observed using a Zeiss LSM 880 laser scanning confocal microscope with 20 x objective. conA_488_ staining (ThermoFisher C11252) was performed as described previously (Habusha et al., 2019). Bacteria were observed using a Zeiss LSM 880 laser scanning confocal microscope with 40 x (water immersion) objective. Fluorescent intensity was measured using ImageJ.

### DNA sequence analysis

DNA was extracted from an overnight culture of the mutant bacterial strain along with the WT strain using DNeasy PowerSoil Pro kit (QIAGEN), according to manufacturer’s instructions. DNA-seq libraries were prepared using Nextra DNA Flex library preparation kit (Illumina), according to manufacturer’s instructions. Illumina NextSeq 500 High-Output 50bp single reads were partially assembled into contigs using the SPAdes algorithm (Bankevich et al., 2012) with default parameters. The contigs were aligned to the genome of the WT bacteria from our lab stock. DNA-seq dataset is available in the SRA repository with the accession numbers PRJNA729435

### Graphs and Statistics

All graphs and statistical tests were performed in Excel. P values throughout the paper were determined using Student t-test, except for figure S3, where a chi-square test was performed.

## Supplementary figures

**Figure S1.**
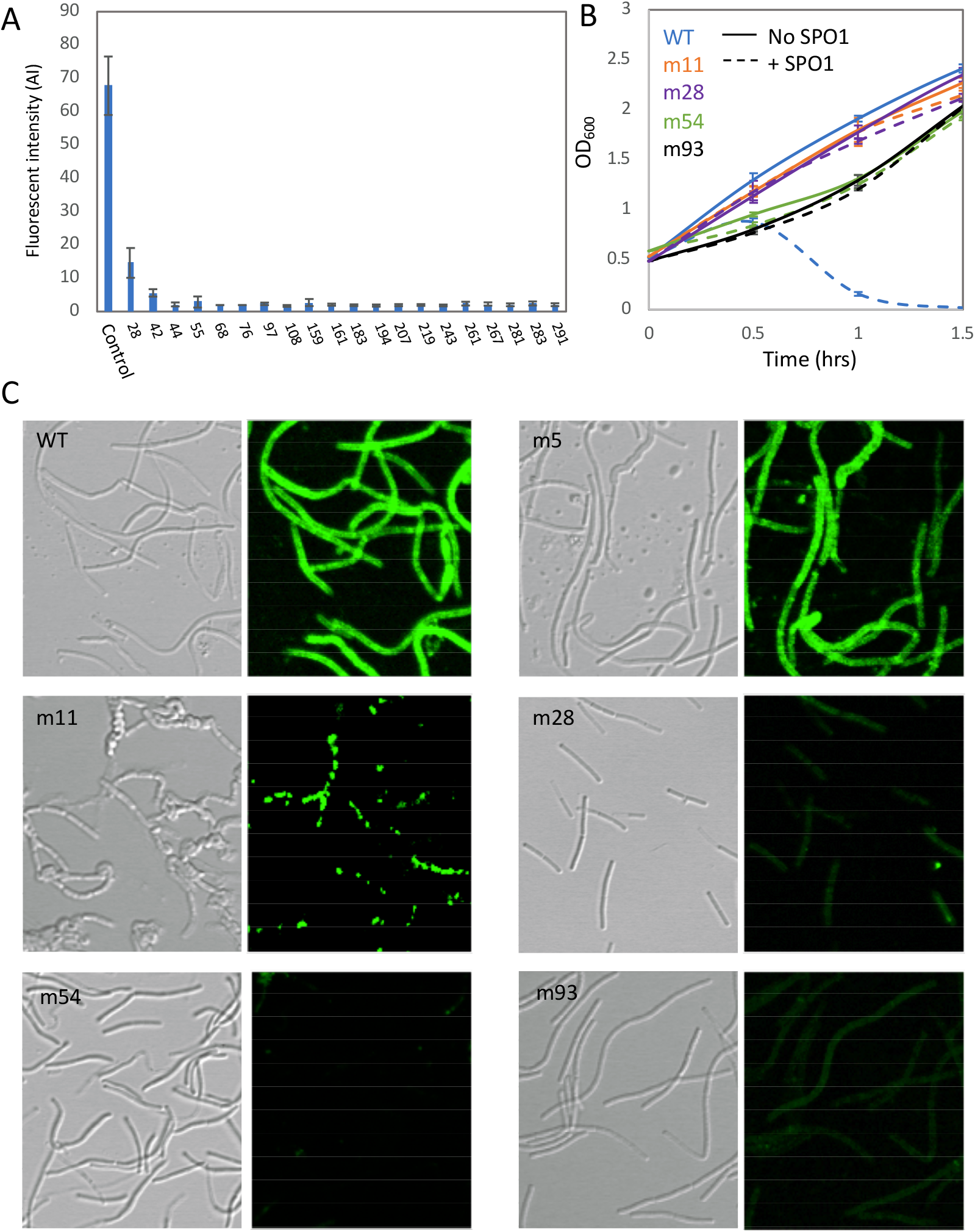
SPO1 resistance profile of phage survival isolates. (**A**) SPO1 resistant bacteria isolated from plants were stained with conA_488_ and observed under the microscope. Shown are average and SD fluorescent intensity values of 10 bacteria from each of the survived colonies. (**B**) The indicated bacterial strains were infected with SPO1 in LB media and OD_600_ was followed. Shown are averages and SD with n = 3. (**C**) The indicated bacterial strains were stained with ConA_488_. Shown are representative DIC (left panels) and fluorescence from ConA_488_ (right panels) images from each strain.

**Figure S2.**
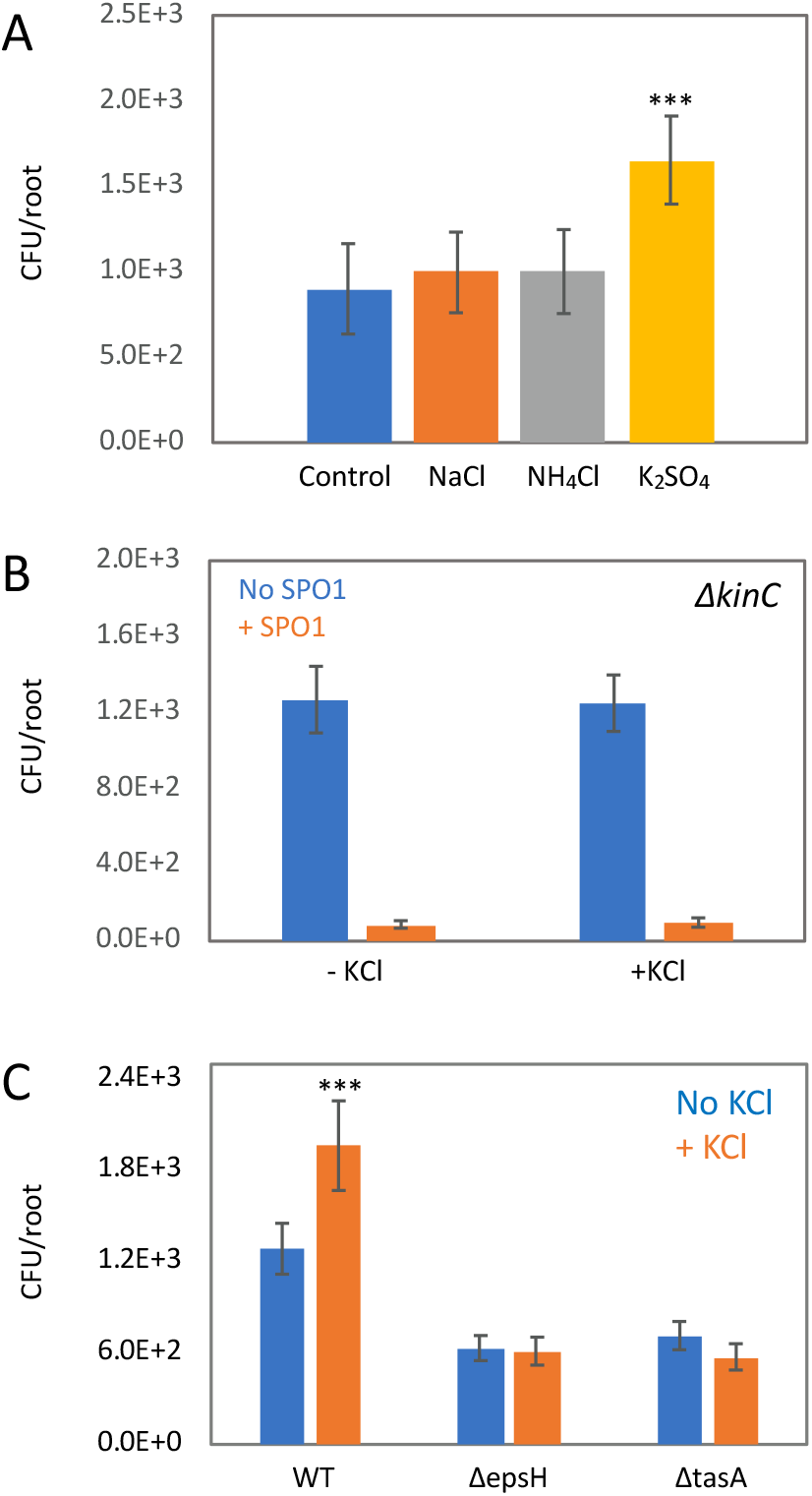
Potassium modulates *B. subtillis* root colonization and phage resistance. (**A**) Seedlings were inoculated with WT bacteria and grown for 48 hrs on 0.25MS agar plates, in the absence or presence of 5mM of the indicated salts, and the number of colonizing bacteria were counted. Shown are averages and SD of 2 independent experiments with n ≥ 3 for each. *** = *P <* 0.005. (**B**) Seedlings were inoculated with Δ*kinC* bacteria, with or without SPO1 addition, and grown for 48 hrs on 0.25MS agar plates, in the presence or absence of 5mM KCl, and the number of colonizing bacteria were counted. Shown are averages and SD of 2 independent experiments with n ≥ 3 for each. (**C**) Seedlings were inoculated with the indicated bacterial strains and grown for 48 hrs on 0.25MS agar plates, in the absence or presence of 5mM KCl, and the number of colonizing bacteria were counted. Shown are averages and SD of 2 independent experiments with n ≥ 3 for each. *** = *P <* 0.005.

**Figure S3.**
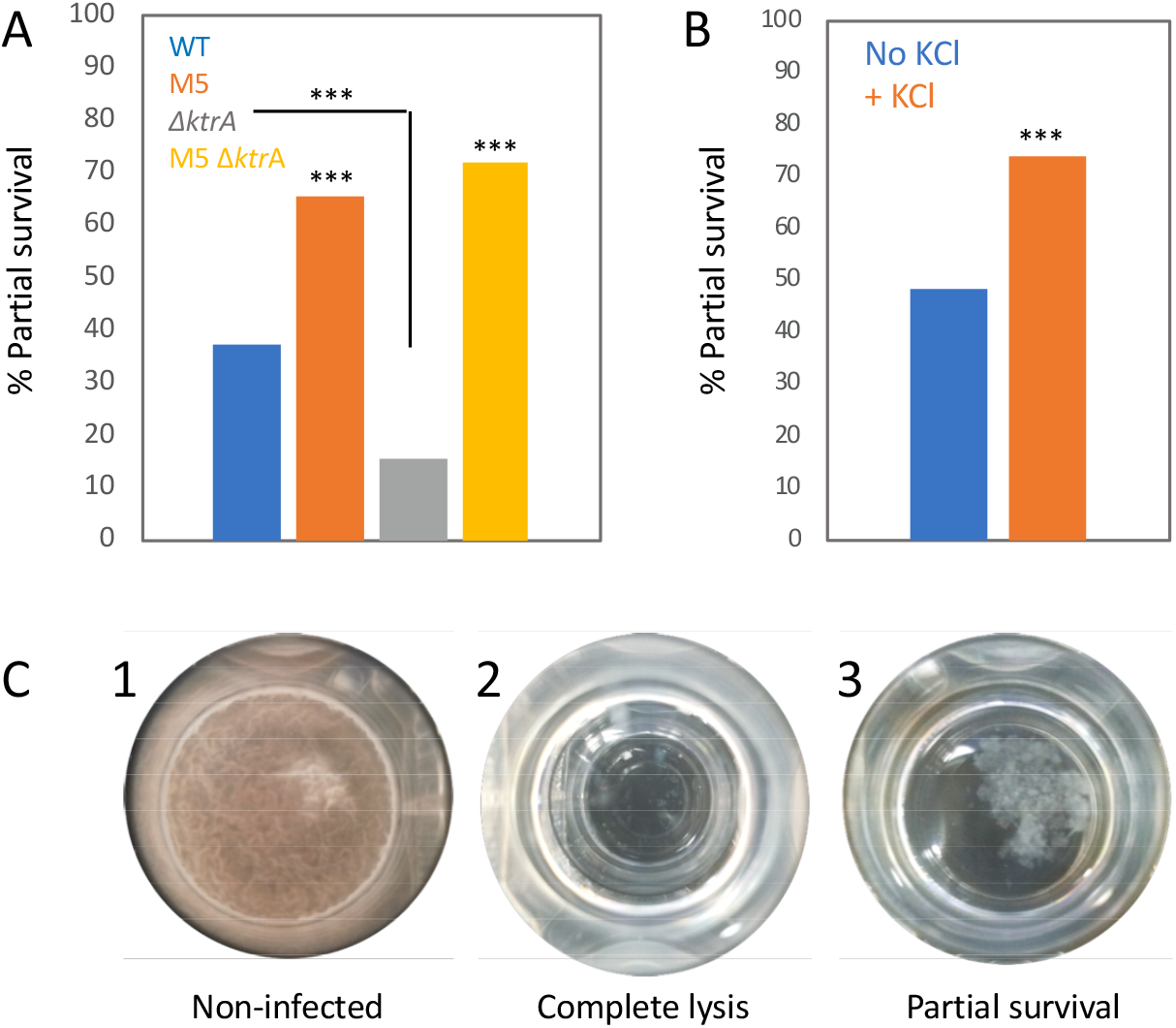
Potassium modulates phage resistance during pellicle biofilm formation. (**A**) The indicated bacterial strains were inoculated into MSgg medium with or without SPO1 addition for 48 hrs, and the number of cultures exhibiting partial SPO1 survival (see C) were counted. Shown is the percentage of the culture surviving SPO1, n ≥ 30. *** = *P <* 0.005. (**B**) WT bacteria were inoculated into MSgg medium with or without 5mM KCl and infected with SPO1 for 48 hrs, as described in A. Shown is the percentage of the culture surviving SPO1, n ≥ 30. *** = *P <* 0.005. (**C**) Shown are representative images for the experiment described in (A and B), for an un-infected culture (1), a culture undergoing complete phage lysis (2) and a culture counted as partially surviving phage infection (3).

